# ReMASTER: Improved phylodynamic simulation for BEAST 2.7

**DOI:** 10.1101/2023.10.09.561485

**Authors:** Timothy G. Vaughan

## Abstract

**Summary:** Phylodynamic models link phylogenetic trees to biologically-relevant parameters such as speciation and extinction rates (macroevolution), effective population sizes and migration rates (ecology and phylogeography), and transmission and removal/recovery rates (epidemiology) to name a few. Being able to simulate phylogenetic trees and population dynamics under these models is the basis for (a) developing and testing of phylodynamic inference algorithms, (b) performing simulation studies which quantify the biases stemming from model-misspecification, and (c) performing so-called model adequacy assessments by simulating samples from the posterior predictive distribution. Here I introduce *ReMASTER*, a package for the BEAST 2 phylogenetic inference platform which provides a simple and efficient approach to specifying and simulating the phylogenetic trees and population dynamics arising from phylodynamic models. ReMASTER is a complete rewrite of an earlier package, MASTER, and boasts improved efficiency, ease of use, flexibility of model specification, and integration with BEAST 2.

**Availability and Implementation:** ReMASTER can be installed directly from the BEAST 2 package manager, and its documentation is available online at https://tgvaughan.github.io/remaster. ReMASTER is free software, and is distributed under version 3 of the GNU General Public License. The Java source code for ReMASTER is available from https://github.com/tgvaughan/remaster.

## 1. Introduction

Stochastic models of population growth and tree generation feature heavily in modern phylogenetics and phylodynamics. This is particularly true in the case of Bayesian inference frameworks, which use parameter-rich generative models to provide prior distributions for phylogenetic trees and, ultimately, to connect genetic sequence data to these dynamics. While not usually a part of the inference procedure itself, direct simulation of the stochastic realisations of these models is useful in many ways. For instance, simulated ensembles are used for independent verification during Markov chain Monte Carlo algorithm development (e.g. [1]). Additionally, simulated trees are often used as the ground truth for simulation studies—investigations into the response of inference algorithms to model misspecification (e.g. [2]). Finally, direct simulation can also be used to produce samples from the posterior predictive distribution as part of phylodynamic model adequacy assessments [3].

Given this central importance of simulation to the field, it is unsurprising that a vast array of purpose-built simulation software exists, and that this is far to extensive to list here in full. For example, in the context of coalescent-based population genetics, tools such as ms [4] and fastsimcoal [5] allow for the extremely rapid simulation of large amounts of genetic data under a wide variety of demographic scenarios for the purpose of inference using approximate Bayesian computation or similar simulation-heavy inference methods. On the other hand, the simulation of linear birth-death phylodynamic models are well-catered for by the R packages TreeSim [6] and Castor [7].

While it is certainly possible to find a simulator capable of producing trees and population histories under almost any given model from among the very large ensemble of existing software, there are two important reasons why it makes sense to provide another tool. Firstly, while simulators for many distinct models exist, very few of these simulators allow for flexible and intuitive specification of most models commonly used in phylodynamics. Specifically, no single system accounts for multi-type models, density dependent effects, and flexible sampling schemes. Secondly, there is great value in having a simulation tool tightly integrated with a phylogenetic inference platform, allowing one to directly and trivially compare simulated and real data, and to construct elaborate joint analyses.

This application note briefly introduces ReMASTER, a flexible phylodynamic simulation tool for the Bayesian phylogenetic inference platform BEAST 2 [8]. It is a complete bottom-up rewrite of a successful earlier package, MASTER [9], which provides a large number of improvements over the original, including:

### More efficient birth-death tree simulation

For birth-death trees, ReMASTER simulates a birth-death trajectory under the corresponding model, then directly simulates the reconstructed phylogeny based on that trajectory. This avoids simulating and keeping in memory the unused parts of the full phylogeny, resulting in a significant reduction in required computation time for birth-death trees. (See Fig. 1.)

**Fig. 1:**
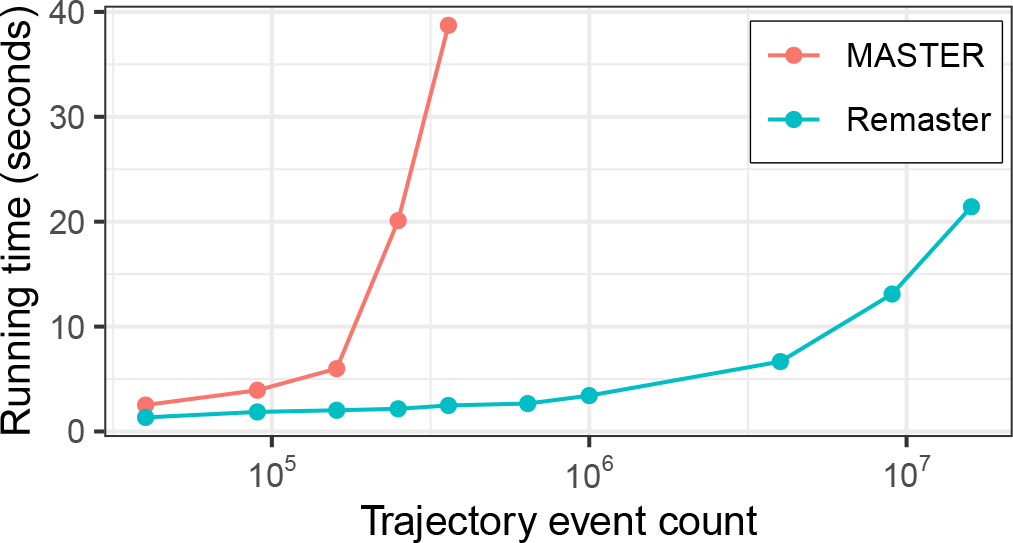
Running times for ReMASTER compared with the older MASTER package, when simulating reconstructed 100-leaf trees under a series of stochastic logistic growth models with increasing carrying capacities. Running times in the figure are shown against the estimated number of simulated birth and death events, which increase quadratically with carrying capacity. MASTER simulates and prunes the full population tree leading to significantly more computation than ReMASTER’s approach of simulating only the reconstructed tree.

### Flexible coalescent simulation

MASTER’s coalescent simulator was very limited, allowing only piecewise-constant variation in effective population sizes. In contrast, ReMASTER allows rates to vary under the influence of any continuously-varying population model implemented in BEAST 2. Further, ReMASTER explicitly supports simulation of trees under coalescent approximations to arbitrary birth-death processes [10].

### Integrated specification of sampling schemes

Like MASTER, ReMASTER expects users to specify birth-death or coalescent models as a series of “reactions”. ReMASTER takes this idea further by incorporating the generation of “samples” (tree leaf nodes) into the same formalism, allowing complex sampling schemes to be much more easily described. This is done by considering all samples to also be the result of a reaction firing, either at different times with an overall (potentially time-dependent) rate, or at fixed times with a specified probability or explicit count. This important extension is discussed further in section 3 below.

### Deeper integration with BEAST 2

MASTER used its own unique data types and mechanisms for specifying trees and model parameters and for producing output files, while ReMASTER uses BEAST 2’s own data types and output logging mechanisms as much as possible. The result of this is that ReMASTER simulation specifications can be easily embedded within standard BEAST 2 analysis scripts, meaning that they can be directly used to initialise BEAST 2’s Markov chain Monte Carlo state or to combine simulation and inference scripts useful for conducting simulation studies or model adequacy assessments.

## 2. Installation and Usage Overview

Installing ReMASTER firstly requires BEAST 2.7, which can be obtained from https://beast2.org. With this in place, ReMASTER can be installed directly using BEAST 2’s bundled package manager. Just like MASTER, using ReMASTER involves firstly constructing an input file describing the phylodynamic model and simulation required. Running BEAST 2 on this input file then runs corresponding simulation, resulting in output files containing the simulated trees and population size trajectories. Tree files use the same NEXUS [11] format used by BEAST 2 phylogenetic inference runs and can be visualised using standard tools such as FigTree [12], IcyTree [13] or ggtree [14]. Population trajectory files can be imported into the R statistical computing environment [15] for further processing or visualisation using a script included with the package.

## 3. Integrated sampling model specification

Here I briefly discuss ReMASTER’s integrated reaction-based approach to specifying birth-death-sampling phylodynamic models.

Specifying a model in a ReMASTER input file firstly involves defining one or more populations, e.g. S, E, I and R for the compartments of a Susceptible-Exposed-Infectious-Removed (SEIR) model, and then specifying one or more reactions determining how these populations change through time.

Reactions determining the eventual dynamics are then specified directly using notation borrowed from stochastic chemical kinetics. For instance, the three main SEIR reactions, corresponding to transmission, becoming infectious, and removal, could be written:

~~~
<reaction spec=“Reaction” rate=“0.1”>
 I + S −> I + E
</reaction>
<reaction spec=“Reaction” rate=“1.0”>
 E −> I
</reaction>
<reaction spec=“Reaction” rate=“0.5”>
 I −> R
</reaction>
~~~

The value of rate in each case is the reaction rate per-reactant configuration, and may vary arbitrarily in a piecewise-constant manner through time.

To simulate trees, additional reactions generating “samples”— corresponding to leaves (or sampled ancestors [16]) in the phylogenetic tree—must be included. In reactions, samples appear simply as individuals belonging to specially marked populations. For instance, the following reaction describes a sampling scheme in which each infectious individual is sampled (but not removed) according to a Poisson process with a rate of 1 sample per time unit after 5 time units. (The rate is zero before this time.)

~~~
<reaction spec=“Reaction” rate=“0 1” change Times=“5”>
 I −> I + sample
</reaction>
~~~

“Punctual” reactions are applied at specified time points, and may be applied either with a particular probability p to every reactant or a pre-specified number of times. While punctual reactions are completely general and don’t necessarily produce samples, they are particularly useful for producing trees with many leaves sampled contemporaneously. For example, the following punctual reaction indicates that each infected individual present at times 5 and 10 should be sampled (and removed) with probability 0.1:

~~~
<reaction spec=“PunctualReaction” p=“0.1” times=“5 10”>
 I −> sample
</reaction>
~~~

With these reactions in the input file, ReMASTER can simulate realisations of both the birth-death population-level trajectories and the phylogenetic trees ancestral to the “samples” generated by the final two reactions above.

While this illustration has focused on birth-death models, leaves in coalescent phylodynamic models are also introduced via reactions. The difference there is that coalescent reactions deal exclusively with lineages ancestral to sampled taxa, so reactions are specified backward in time, and no special “sample” population type is necessary to generate leaves.

Making sample-production an explicit part of the model in this way allows for succinct and intuitive specification of a very wide range of phylodynamic scenarios.

## 4. Closing remarks

Full installation and usage instructions, together with a comprehensive manual describing the model description language, output processing guidelines and example input files, are available at https://tgvaughan.github.io/remaster.

ReMASTER represents a dramatic improvement in usability, flexibility and efficiency over MASTER, and I strongly recommend its use over the earlier package for all future work.

## 6. Author contributions statement

TGV wrote the software and the manuscript.

## 7. Acknowledgments

The author was supported by ETH Zürich.

